# The genomic architecture of adaptation to larval malnutrition points to a trade-off with adult starvation resistance in *Drosophila*

**DOI:** 10.1101/2020.12.01.406686

**Authors:** Tadeusz J. Kawecki, Berra Erkosar, Cindy Dupuis, Brian Hollis, R. Craig Stillwell, Martin Kapun

## Abstract

Periods of nutrient shortage impose strong selection on animal populations. Experimental studies of genetic adaptation to nutrient shortage largely focus on resistance to acute starvation at adult stage; it is not clear how conclusions drawn from these studies extrapolate to other forms of nutritional stress. We studied the genomic signature of adaptation to chronic juvenile malnutrition in six populations *of Drosophila melanogaster* evolved for 150 generations on an extremely nutrient-poor larval diet. Comparison with control populations evolved on standard food revealed repeatable genomic differentiation between the two set of population, involving >3,000 candidate SNPs forming >100 independently evolving clusters. The candidate genomic regions were enriched in genes implicated in hormone, carbohydrate, and lipid metabolism, including some with known effects on fitness-related life-history traits. Rather than being close to fixation, a substantial fraction of candidate SNPs segregated at intermediate allele frequencies in all malnutrition-adapted populations. This, together with patterns of among-population variation in allele frequencies and estimates of Tajima’s *D*, suggests that the poor diet results in balancing selection on some genomic regions. Our candidate genes for tolerance to larval malnutrition showed a high overlap with genes previously implicated in acute starvation resistance. However, adaptation to larval malnutrition in our study was associated with reduced tolerance to acute adult starvation. Thus, rather than reflecting synergy, the shared genomic architecture appears to mediate an evolutionary trade-off between tolerances to these two forms of nutritional stress.

## Introduction

The availability and quality of organic nutrients is a key ecological factor that determines the survival and fitness of animals. Natural selection imposed by periods of nutrient shortage is therefore thought to have shaped manifold aspects of animal physiology, life history and behavior (Baker *et al*., 2004; Boggs & Freeman, 2005; Behrman *et al*., 2015; McNamara *et al*., 2016). Resulting adaptations may impart many aspects of performance such as susceptibility to metabolic or infectious disease (Vijendravarma *et al*., 2015; Hardy *et al*., 2018), and thus be relevant for human health (Prentice, 2005; Wells, 2006).

Most of research investigating the phenotypic and genetic bases of evolutionary adaptation to nutritional stress focuses on acute starvation resistance, i.e., the ability to survive periods of complete food deprivation, mainly at adult stage; much of this work has been done in *Drosophila.* Selection for greater starvation resistance of adult fruit flies results in longer development time, greater longevity, increased lipid storage, larger body size and reduced fecundity, which suggests physiological trade-offs between starvation resistance and other fitness-related traits (reviewed in Rion & Kawecki, 2007; Kubrak *et al*., 2017; Hardy *et al*., 2018; Michalak *et al*., 2018). This implies that frequent exposure to starvation favors greater energy reserves, achieved by increased lipid or carbohydrate storage and larger body size (Rion & Kawecki, 2007; Kubrak *et al*., 2017). Furthermore, food-deprived animals appear to switch to a “survival mode” to withstand short periods of malnutrition by reducing metabolic processes and diverting available resources from reproduction (Tatar *et al*., 2003; Rion & Kawecki, 2007). Evolve and resequence studies, which combine experimental evolution and genomic analyses (Turner *et al*., 2011; Kofler & Schlötterer, 2014), indicate that the genomic architecture of acute starvation resistance is complex and highly polygenic, with candidate SNPs in genes affecting lifespan, feeding behavior, catabolic metabolism and lipid body structure and function (Hardy *et al*., 2018; Michalak *et al*., 2018).

In contrast to acute starvation, we know much less about the physiological and genomic bases of adaptation to prolonged nutrient shortage – in particular at the juvenile stage. Juveniles are often the first to suffer from food shortage (Ronget *et al*., 2017), and periods of juvenile malnutrition often have long-lasting consequences for adult fitness (Lindström, 1999; Wells, 2007; Koyama & Mirth, 2018). Juveniles will have little chance to accumulate any metabolic reserve if exposed to nutritional shortage from birth; they must not only survive, but also develop and grow with whatever nutrients they manage to acquire. Waiting out until better times by relying on energy reserves, which is a major target of selection for acute starvation resistance, is not a viable option under chronic juvenile malnutrition. Adaptation to chronic juvenile undernutrition is therefore expected to involve at least partly different mechanisms than resistance to acute starvation. Consistent with this, while adult *Drosophila* selected for starvation resistance have larger body weight with higher lipid stores (Rion & Kawecki, 2007), those selected for tolerance to poor larval diet evolve smaller adult body size without enhanced lipid content (Kolss *et al*., 2009; Vijendravarma *et al*., 2012a). Thus, we hypothesized that adaptation to chronic juvenile malnutrition may involve genes and molecular mechanisms largely different from those mediating adult starvation resistance.

In this paper we investigate the genomic architecture of adaptation to chronic larval undernutrition in six replicated populations of *Drosophila melanogaster* subject to 150 generations of experimental evolution on a nutrient-poor larval diet. The poor diet imposes a strong nutritional stress: larvae from non-adapted populations suffer high mortality and the survivors take twice as long to develop until pupation on the poor diet than on standard diet; yet, the adults still emerge at half of the normal size (Kolss *et al*., 2009; Erkosar *et al*., 2017). Tolerance to the poor diet evolved by the experimental populations manifests in improved survival, faster larval growth and faster development (Kolss *et al*., 2009). It has been associated with changes in larval behavior (Vijendravarma *et al*., 2012b; Narasimha *et al*., 2015) and in digestive physiology (notably with increased activity of digestive proteases; Erkosar *et al*., 2017), but also with increased susceptibility to an intestinal bacterial pathogen (Vijendravarma *et al.*, 2015).

We use whole genome pooled sequencing to analyze the architecture of genomic divergence between the six populations evolved on the poor larval diet (‘Selected’ populations) and the six ‘Control’ populations derived from the same gene pool but evolved on standard diet. We identify and annotate candidate SNPs putatively underlying the adaptive divergence and find that they show contrasting patterns of allele frequencies suggestive of being driven by different modes of selection. We also combine the genome sequence results with differences in gene expression patterns (previously characterized by Erkosar *et al*., 2017).

We then address the evolutionary relationship between adaptation to larval malnutrition and resistance to adult starvation. In contrast to our expectation expressed above, we find that experimental evolution in response to these two forms of nutritional stress involves overlapping sets of genes. Yet, rather than a common physiological mechanism, the shared genetic architecture appears to mediate a trade-off between tolerance to larval malnutrition and resistance to adult starvation.

## Results

### Genome-wide patterns of variation and differentiation

We sequenced pools of 400 females from each of the six Selected populations (evolved on poor diet) and six Control populations (maintained on standard diet). We identified 976,247 high confidence single-nucleotide polymorphic sites (SNPs) that satisfied stringent SNP calling criteria. The SNPs were spread across the five major chromosomal arms (X, 2L, 2R, 3L, 3R), with a few on the dot chromosome 4. Principal component analysis on allele frequencies of all these SNPs clearly separated the Selected from the Control populations, mainly along the first principal component axis, which accounted for 19% of the SNP variation (Figure 1, Welch’s t-test on the first principal component scores *t*_7_ = 13.1, *P* < 0.0001). These results indicate that 150 generations of evolution on the poor versus standard diet resulted in a replicable differentiation of gene pools, with adaptation to the poor diet being to a large degree parallel at the genomic level across the replicate Selected populations.

**Figure 1.**
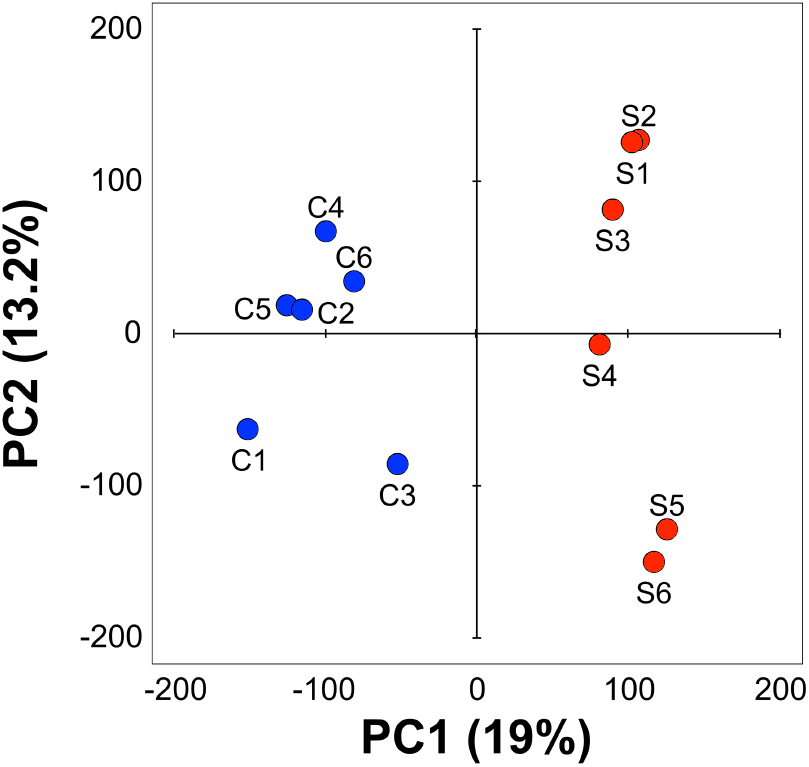
Principal component analysis on all SNPs. The Selected (red) and Control (blue) populations plotted on the first two principal component axes derived from PCA on allele frequencies at all 976,247 SNPs that had at least 10-fold coverage in each population. The % values refer to the fraction of total variance explained.

Despite the same census population sizes, selection under the harsh conditions of the poor diet was expected to reduce the effective population size *(N*_e_) of Selected populations relative to Controls by increasing variance in reproductive success (Barton, 2000). To assess to what degree this was the case, we estimated *N*_e_ based on among-population variance in allele frequencies at putatively neutral SNPs (synonymous and in short introns). Although these estimates did tend to be smaller for Selected than Control populations, the difference was small: 120 versus 144 (*P* = 0.15) for the two main autosomes (chromosomes 2 and 3), 65 versus 74 for the X chromosome (*P* = 0.59, randomization test, see Methods). Interestingly, *N_e_* estimates for the X chromosome are about half of those of autosomes; the theoretical expectation is that *N_e_* of the X should be ¾ of *N_e_* of autosomes (Hartl & Clark, 1997). Genome-wide patterns of within-population variation (quantified as nucleotide diversity *π* and Watterson’s 0) were likewise similar between the evolutionary regimes (see Supporting Information, Supplementary Figure S1 and Supplementary Table S1). Thus, although evolution on the poor versus standard diet resulted in clear differentiation of the gene pools, differences between regimes in within-population patterns of genetic variation were at most minor.

For each replicate population, we further estimated the population genetic statistic Tajima’s *D* for non-overlapping genomic windows of 200 kbp length and on a genome-wide scale. *D* summarizes the shape of the site frequency spectrum and provides information about potential adaptive and non-adaptive evolutionary forces at play (Nielsen, 2005). Negative Tajima’s *D* indicates an excess of low-frequency variants which may be the result of selective sweeps or population size expansion, whereas positive values of *D* imply an excess of intermediate allele frequencies, for example, caused by balancing selection or reductions in population size. Our analyses revealed that genome-wide Tajima’s *D* was overall positive and significantly different from zero (one-sample Student’s t-test) in all populations (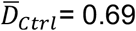, SD = 0.09, *P* < 0.001; 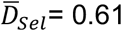, SD = 0.08, *P* < 0.001) without significant differences among the regimes (Mann-Whitney U test; *P* = 0.25). This is consistent with either an initial loss of genetic variation when setting up the experimental populations or with reductions of effective population size in response to the evolutionary regime both in the Selected and the Control populations. Variation in Tajima’s *D* among genomic regions (see Figure 2A) may indicate different types of selection in action. Besides directional selection in the Selected Population, these potentially also include balancing selection resulting in regional positive Tajima’s *D* in the Selected populations, or directional selection in the Control populations (see below).

**Figure 2.**
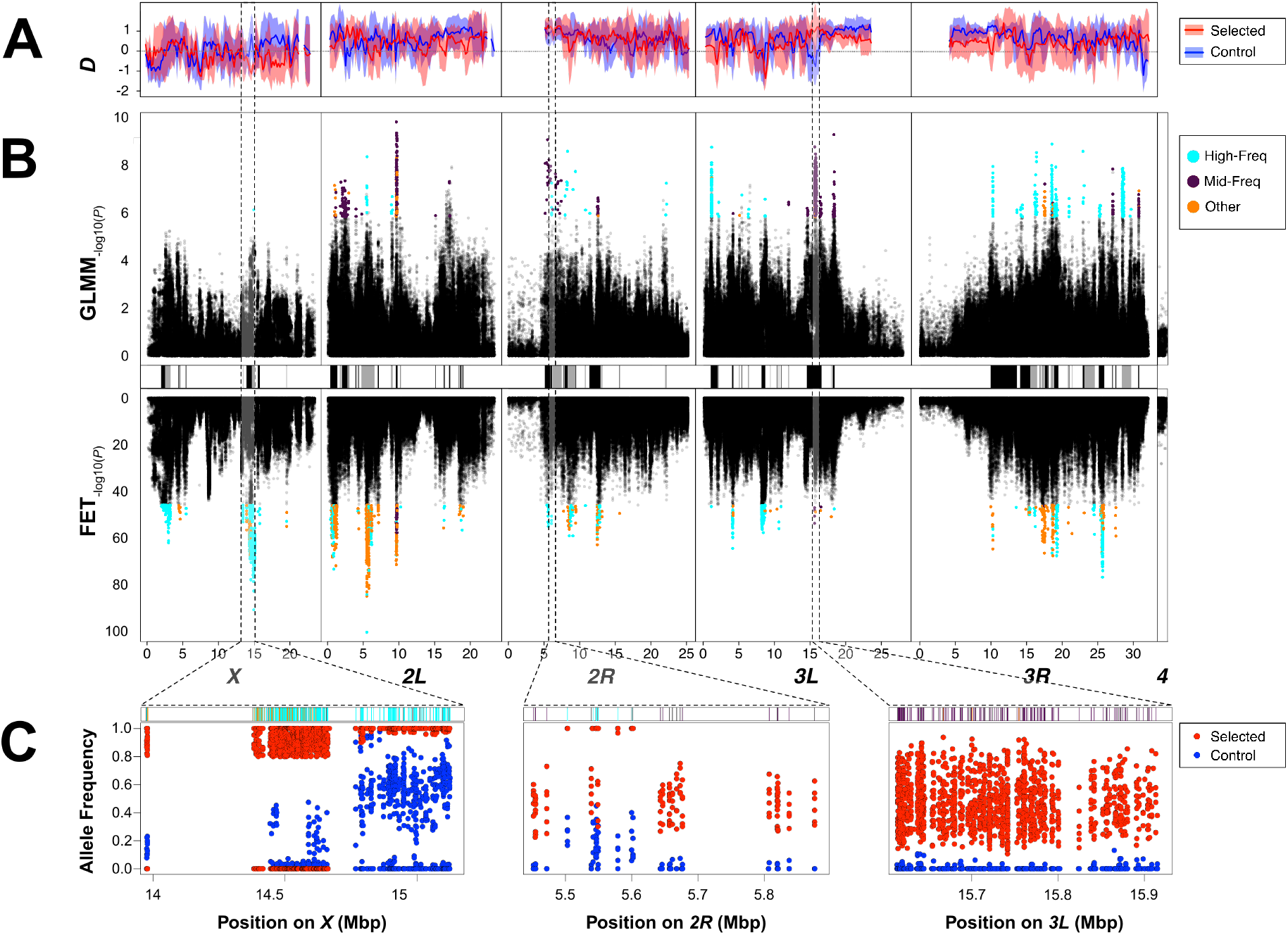
Population genetic analysis of candidates for selection. (A) The top panel show chromosome-wise averages for the population genetic estimator Tajima’s *D* in nonoverlapping windows of 200 kbp size. Solid lines and semi-transparent polygons show means and standard deviations for these estimators, respectively, that were calculated from six selected (red) and six control (blue) replicate populations. (B) The central figures shows Manhattan plots depicting SNP-wise -log_10_ – transformed nominal *P*-values from generalized linear models (GLMM; top panel) and Fisher Exact tests (FET; bottom panel). Putative candidates for balancing selection are highlighted in orange, candidates for directional selection in cyan and candidates with uncertain selection type are shown in dark purple. The black and grey bars separating the upper and lower Manhattan plots indicate linkage blocks which were inferred by correlation of allele frequencies among neighboring SNPs (C) The bottom figures show the distribution of allele frequencies for candidate SNPs in six selected (red dots) and six control (blue dots) populations focusing on three genomic regions that harbor clusters of candidates SNPs potentially affected by balancing or directional selection. All candidate SNPs were conditioned for the alleles that were on average more frequent in the selected populations.

### Candidate SNPs form many independently evolving clusters

To identify candidate SNPs underlying differentiation between the Selected and Control populations we employed a statistical approach based on combining Fisher’s exact tests (FET) and generalized linear mixed models (GLMM). The GLMM emphasizes parallel changes in all replicate populations whereas the FET-based approach emphasizes the difference in mean allele frequencies and can detect cases where an allele becomes fixed in most replicate populations but lost by drift in one. The false discovery rate (FDR) was estimated based on permutations following Jha et al. (2015a; for details and rationale see Methods). From the SNPs that passed the FDR = 0.05 threshold for at least one of the tests we focused on SNPs at which the average allele frequency difference between regimes was greater than the arbitrary threshold of 0.3. (For details and rationale of this approach see Methods.) This approach yielded 2,483 and 1,140 candidate SNPs from FETs and GLMMs, respectively, resulting in a total of 3,425 candidate SNPs (only 97 of those SNPs were shared between both sets; see Supplementary Table S2 and Figure 2B; the color-coded categories of SNPs are defined in the next section).

The candidate SNPs formed clusters characterized by similar patterns of allele frequencies (Figure 2B,C), consistent with selection on a relatively small number of target SNPs and genetic hitchhiking of neighboring SNPs at linkage disequilibrium with the target SNPs. Polymorphic sites that are constrained to evolve together as a consequence of linkage disequilibrium should vary in a highly correlated manner among replicate populations. Therefore, to assess the degree of hitchhiking, we estimated within-regime correlations of allele frequencies between pairs of candidate SNPs (i.e., correlations of residuals from regime means across the 12 populations) *r_w_*. The imprint of linkage disequilibrium is visible as a large excess of *r_w_* > 0.8 between candidate SNPs on the same chromosomal arm, relative to the distribution of *r_w_* between pairs of SNPs on different chromosomes (Figure 3A). Analysis of *r_w_* indicated the existence of massive blocks of highly correlated SNPs on chromosomes *X* and *3L* (Figure 3B, above diagonal). The SNPs forming these blocks are closely linked, as indicated by low pairwise recombination rates (Figure 3B, below diagonal). However, there are other parts of the genome – notably on chromosomal arm 2R, but also on 2L and 3R – that show at most small correlated blocks, suggesting that in those regions many candidate SNPs were free to evolve independently, despite similarly low recombination rates. Thus, the size of co-evolving candidate SNP clusters is not determined by recombination rates alone.

**Figure 3.**
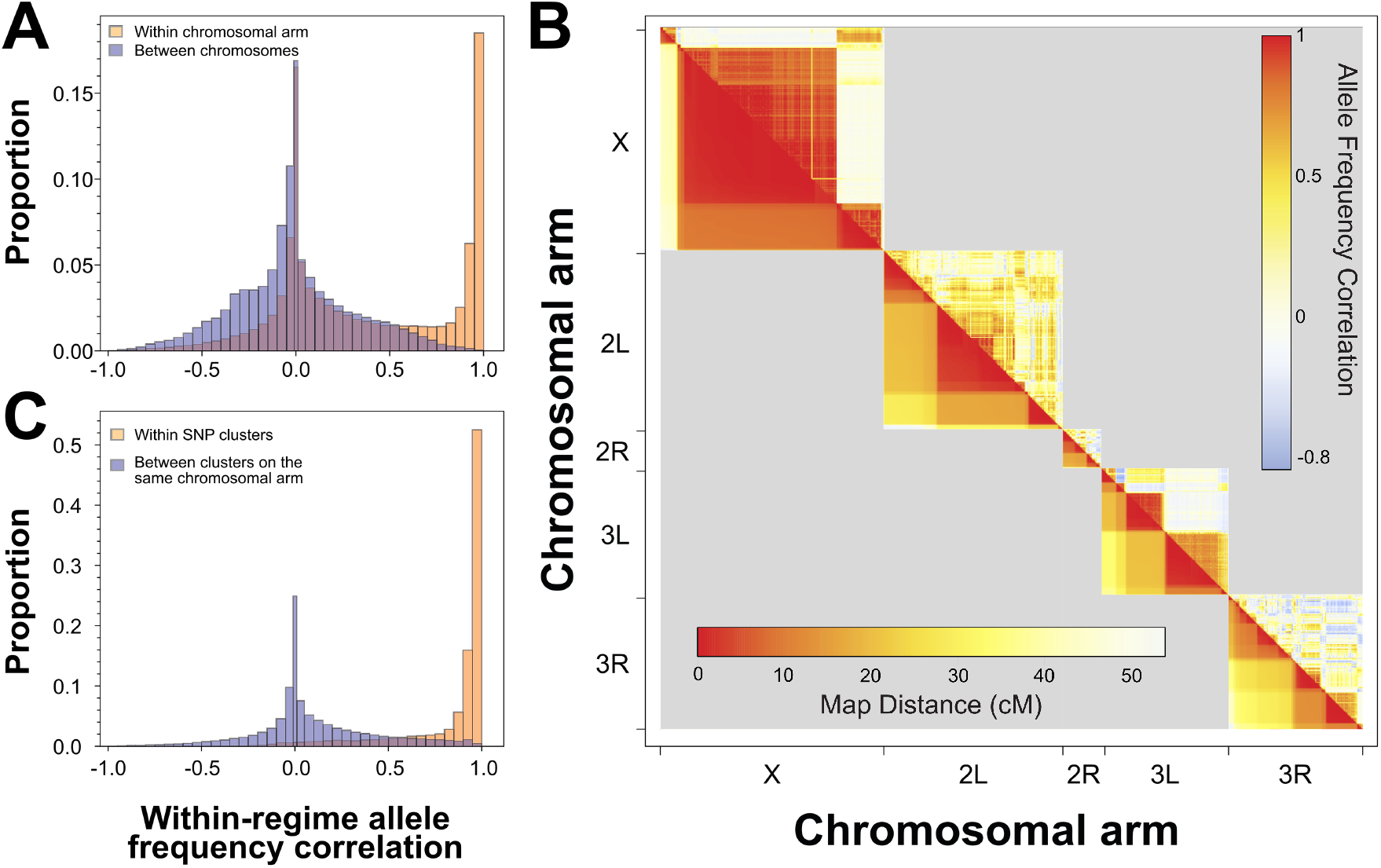
Correlations of allele frequencies between candidate SNPs indicate clusters of SNPs at linkage disequilibrium. (A) Frequency distribution of the within-regime correlations between candidate SNPs on the same chromosomal arm (yellow) compared to the corresponding distribution for pairs of SNPs on different chromosomes (purple). (B) Above diagonal: Heat map of within-regime correlations of the frequency of the “selected” allele between candidate SNPs on the same chromosomal arm. The heat values are Pearson’s correlation coefficients of residuals of population allele frequencies from their corresponding regime means, calculated for each pair of SNPs across the 12 populations. Below diagonal: pairwise recombination rates (cM) between candidate SNPs on the same chromosomal arm based on estimates from Comeron *et al.* (2012). The axes preserve the order of SNPs on the chromosome, but not the distance between them; thus, the size of clusters of correlated SNPs in the plot is proportional to the number of candidate SNPs in the cluster but not to the spatial extent of the cluster on the chromosome. (C) Frequency distribution of the within-regime correlations between SNPs assigned to the same cluster (yellow) and between SNPs sharing chromosomal arm but assigned to different clusters (purple); for criteria defining the clusters see text.

We used the within-regime correlation *r_w_* to delineate clusters of linked candidates SNPs whose evolution was apparently strongly bound together by linkage disequilibrium. Two neighboring candidate SNPs were included in the same cluster if (1) distance between them was less than 200 kb and (2) any SNP within 200 kb left of the midpoint between them was correlated with any SNP within 200 kb right of the midpoint with *r_w_* > 0.8. While necessarily arbitrary, this criterion was effective in grouping most highly correlated SNPs within the same cluster, with correlations between clusters on the same chromosomal arm having a similar distribution to correlation of SNPs on different chromosomes (Figure 3C). Of 131 putative candidate SNP clusters defined by this approach, 48 consisted of a single SNP. At the other end of the spectrum, a cluster on the X chromosome contained 744 candidate SNPs within a 685 kb block, and the largest cluster (on chromosome 3R) spanned 3.5 Mb although it only contained 26 candidate SNPs (Supplementary Table S3).

Such large linkage blocks may result from chromosomal inversions, which suppress recombination in heterozygous state (Hoffmann & Rieseberg, 2008; Kirkpatrick, 2010; Kapun & Flatt, 2018). However, an indirect analytical approach (Kapun *et al*., 2014) indicated that the known chromosomal inversions were absent or at very low frequencies (< 0.05) in all replicate populations and thus unlikely to have contributed to the evolutionary response in our experiment (see additional analyses in the Supporting Information).

### Distinct allele frequency patterns at candidate SNPs

Patterns of allele frequencies at candidate SNP loci can be suggestive of the mode of selection driving them (Nielsen, 2005). By convention, in the following we refer to the alleles with higher mean frequency in the Selected than in the Control populations as the “selected” alleles. For about 27% of candidate SNPs the “selected” alleles were fixed or close to fixation (frequency > 0.9) in all Selected populations while showing intermediate to low frequencies in Controls, typically quite variable among replicate Control populations (e.g., the region around position X: 15MB, Figure 2C left). This pattern is consistent with directional selection in the Selected populations (i.e., on the poor diet) and weak or no selection in the Controls. At another 17% of candidate SNPs the “selected” allele was fixed or at a high frequency at five Selected population but absent from the sixth; almost all of these SNPs were within an apparent large linkage block on chromosome X between positions 14.37 and 14.66 Mb, where the Selected population S4 lacked the “selected” allele (Figure 2C left). This pattern is also consistent with directional selection on some of the SNPs in the cluster, but with the selected haplotype being lost due to drift/founder effect in one of the Selected populations.

In contrast, many other candidate SNPs showed a very different pattern, being polymorphic with intermediate frequencies in all Selected populations, with the “selected” allele being lost or at low frequencies in Controls (e.g., the region around positions 3L: 15.5-15.8 Mb, Figure 2C right). While this pattern could occur under other scenarios (see Discussion), it is consistent with balancing selection favoring initially rare alleles and subsequently maintaining them an intermediate frequency in Selected populations. Balancing selection would not only act to maintain polymorphism, but also tend to keep allele frequencies close to a polymorphic equilibrium and hence reduce variation among replicate populations evolving under the same regime. Thus, if balancing selection played a greater role in the Selected than in the Control populations then among-population variance in allele frequency at candidate SNPs should tend to be smaller in the Selected than in the Controls.

To address this prediction, we calculated the standard deviation of population allele frequencies at each candidate SNP for each regime separately. We then compared these standard deviations between Selected and Control populations, binned by the respective regime means. Candidate SNPs with mean frequency in Selected populations in the 0.25-0.75 range were generally characterized by lower among-population variance than candidate SNPs satisfying the corresponding criterion in Control populations (Figure 4). To formally test for this difference while accounting for the non-independence of linked SNPs, we relied on the 131 putatively independent candidate SNP clusters defined above. For each cluster we picked the SNP whose mean allele frequency in Selected populations was closest to 0.5, and of those we retained 28 SNPs with the mean allele frequency in the 0.4-0.6 range. Analogous procedure identified 23 SNPs from different clusters whose mean frequency in Control populations was in the 0.4-0.6 range. Comparison of these two small sets of SNPs confirmed that SNPs with intermediate mean frequency in Selected populations were less variable among replicate Selected populations than the SNPs with intermediate mean frequency in Controls were variable among replicate Control populations (median SD 0.15 versus 0.22, *P* = 0.011, Mann-Whitney U Test).

**Figure 4.**
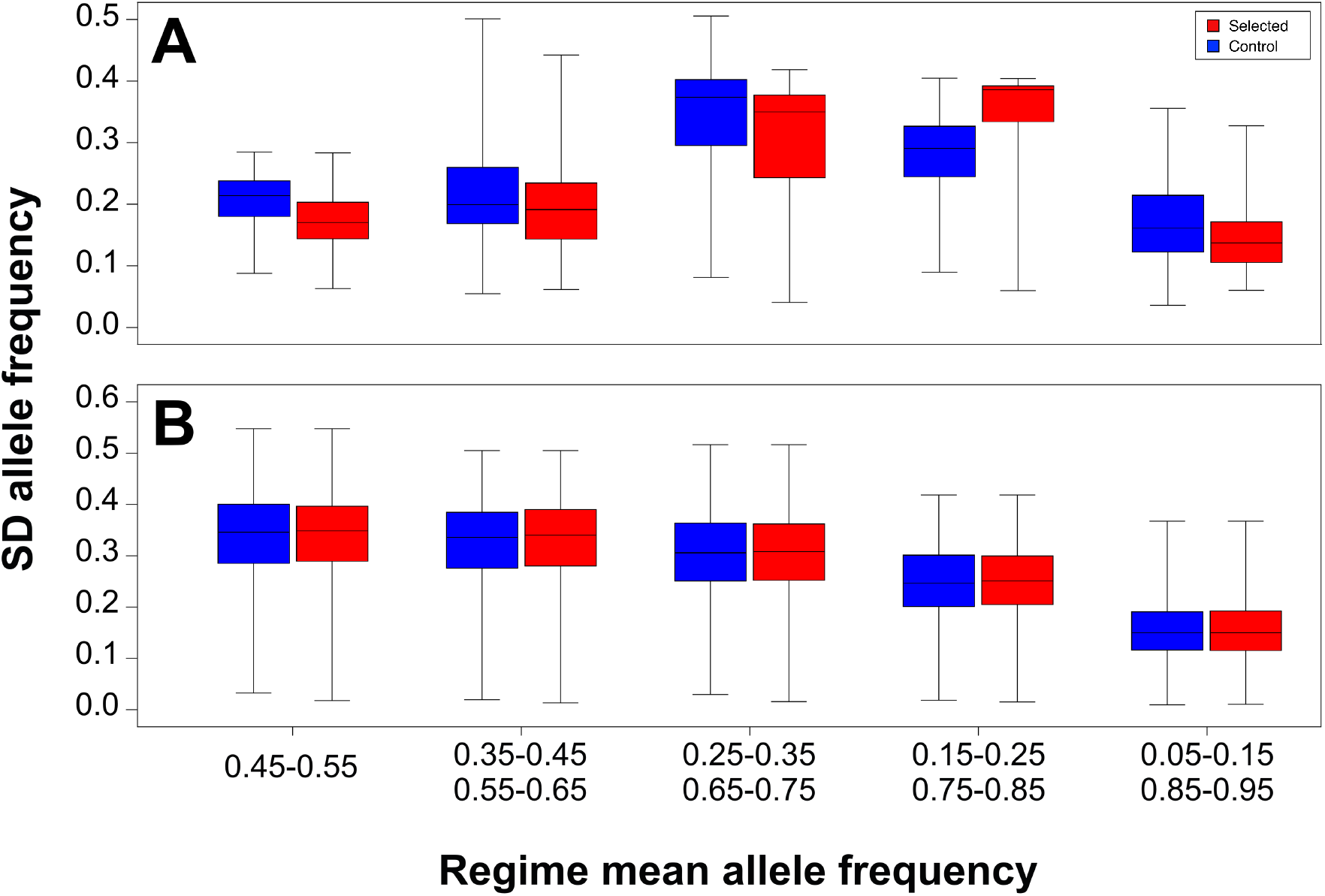
Variation in allele frequency at candidate SNPs among replicate Selected and Control populations. Standard deviation of allele frequency calculated separately for each SNP among Selected (red) and Control (blue) populations and binned according to the mean frequency of the SNP in the respective regimes based on (A) candidate SNPs and (B) randomly drawn non-candidate SNPs.

No such difference in among-population variation was observed for non-candidate SNPs (Figure 4B), for which the median variance for Selected populations was slightly higher than for Controls, in agreement with the lower *N_e_* estimates of the former. Thus, the lower among-population variation in allele frequency in the Selected than Control populations at SNPs that are far from fixation is specific to candidate SNPs. It should be noted that the statistical tests used to detect the candidate SNPs, as well as the above comparison of variation, are both symmetric with respect to evolutionary regimes and the identity of the alleles. Therefore, this difference in the pattern of variance of candidate SNPs suggests that diet-specific selection acts to maintain polymorphism to a greater degree in the Selected than in the Control populations.

In order to see if candidate SNPs showing these two distinct allele frequency patterns show different functional annotations, we operationally defined two subsets of candidates. “High frequency” candidates were defined as those with frequencies of the “selected” allele ≥ 0.75 in all Selected populations, or in five Selected populations and being lost from the sixth population (1,818 SNPs or 53 % of candidates, cyan in Figure 2B). “Mid-frequency” candidates were defined as candidate SNPs at which frequencies of the “selected” allele in each replicate Selected population ranged between 0.1 and 0.9 and whose mean allele frequency in the Selected populations was between 0.25 and 0.75 (548 SNPs 16% of candidates, spread across all autosomal arms; purple in Figure 2B). As can be seen in Figure 2B, these two types of SNPs mainly occurred in different clusters. The remaining 1,059 candidate SNPs did not correspond to either of the above criteria (orange in Figure 2B). SNP classified to these different classes are color-coded in Figure 2B.

### Candidates are enriched for metabolic and hormonal signaling genes

Only 124 candidate SNPs (3.6 %) in 72 genes were annotated as non-synonymous coding (Supplementary Table S2), roughly what would be expected based on the total number of non-synonymous SNPs. Relative to an expectation based on genomic distribution of non-candidate SNPs, we found a significant excess of candidates in intergenic regions and a deficiency in regions comprising 2 kb up- and downstream from genes (Supplementary Figure S2). However, we found a significant excess of candidate SNPs in known regulatory regions (REDfly database; Gallo *et al*., 2006).

Using SNPeff (v.4.2; Cingolani *et al*., 2012) we annotated the 2,511 candidate SNPs to 771 genes (many SNPs were annotated to more than one gene) and performed a GO-term enrichment analysis using Gowinda (Kofler & Schlötterer, 2012). Genes associated with candidate SNPs showed a significant enrichment of GO categories related to hormonal signaling and metabolic processes (Supplementary Table S4). They include key genes in ecdysteroid metabolism, notably *ecdysone oxidase (Eo),* with three missense candidate SNPs in its coding region, and several of its paralogs (fiz, CG9512, CG9503, CG9514, CG12398; Takeuchi *et al*., 2005), as well as *dib* involved in early stages of ecdysteroid synthesis (Warren *et al*., 2002). Three transcription factors that play a key role in triggering metamorphosis in response to ecdysone *(Eip75B, Kr-h1* and *fkh)* are likewise associated with candidate SNPs. Also associated with candidate SNPs are key members of insulin receptor signaling, including *foxo*, a major transcription factor mediating transcriptomic response to reduced insulin signaling (Alic *et al*., 2011), *dock,* which is a negative regulator of insulin signaling (Willoughby *et al*., 2017), and the fat-body expressed *ilp6,* which regulates metabolic response to nutrient shortage (Chatterjee *et al*., 2014). Interestingly – given the context of chronic nutrient shortage – we found no indication of the nutrient-sensing TOR signaling to have been targeted; none of the 62 genes in the GO term “TOR signaling” was associated with a candidate SNP. Among genes involved in metabolism, multiple genes directly involved in accumulation of triglycerides for storage *(SCAP, Lpin, Acsl, Dgat-2)* and their mobilization for catabolism (Lsd-2, dob and pdgy; Heier & Kühnlein, 2018) are associated with candidate SNPs, in particular of those showing the “high frequency” pattern described above (Supplementary Table S4).

Significantly enriched GO terms (Supplementary Table S4) also include a category of glutamate receptors involved in neuromuscular junctions in the larval body wall, one *(GlurIIB)* with two missense SNPs, and salivary gland proteins involved in adhesion of pupae to surfaces, again with one glue gene *(Sgs4)* with missense SNPs and one (*chc*) previously found to have a lower expression in Selected larvae (Erkosar *et al*., 2017). While these may seem idiosyncratic, they can be linked to phenotypic differences between the Selected and Control populations (see Discussion).

Interestingly, when testing for enrichment separately in the “high frequency” and “midfrequency” sets of candidate genes defined in the preceding section, we found no overlap in significantly enriched GO terms. As in the analysis of all candidate SNPs, we found that the “high frequency” candidates were enriched in genes involved in hormonal metabolic processes, lipid metabolic processes and Insulin/Insulin-like signaling, whereas the “mid-frequency” candidates were mainly associated with sugar/amino-sugar transferases (e.g. *sff, sxc, GlcAT-S).* The statistical significance of top GO term enrichment was markedly higher in this separate analysis of “high frequency” and “mid-frequency” types than when all candidate SNP were analyzed jointly (Table 1). This suggests that these two categories of candidate SNPs not only show different allele frequency patterns, but are also functionally different.

### Functional links between genomic and transcriptomic candidate genes

The excess of candidate SNPs in known regulatory regions suggests that the dietary adaptation has been at least partly mediated by regulatory changes, consistent with previously reported differences in gene expression patterns between the Selected and Control populations (Erkosar *et al*., 2017). To explore the degree to which such changes may have been cis-regulatory, we assessed the overlap between genes identified as differentially expressed by Erkosar et al. (2017) and genes to which candidate SNPs from this study were annotated. We found 102 such candidates showing both SNP and expression differentiation (Supplementary Figure S3). However, while the overlap between the SNP and expression candidates is not greater than expected by chance (*P* = 0.33, Super Exact Test), these candidates include numerous genes involved in insulin signaling and lipid metabolism already mentioned in the previous section: *foxo, Ilp6, dock, Acsl, Lpin, Lsd-2, dob, pdgy;* all are downregulated in the Selected populations except for *dock*– which is a negative regulator of insulin signaling. It also includes *fiz,* a putative ecdysone oxidase with natural variants that affect larval growth (Glaser-Schmitt & Parsch, 2018), which is strongly downregulated in Selected populations and has several candidate SNPs closely upstream from the coding region. Thus, although the list of potential candidates for cis-regulatory evolution is short, it includes a number of genes whose function is highly relevant to response to nutrition.

To explore potential functional links between genomic and expression candidates, and to see to what degree the different candidate genes interact physically, we constructed networks of candidate genes based on the existence of known interactions between them (based on the DroID database, (Yu *et al*., 2008; Murali *et al*., 2011)). The combined set of all genomic and transcriptomic candidate genes was characterized by 2.3 pairwise interactions per gene on average, about 40% more than expected based on a randomly drawn set of genes of the same size (*P* = 0.011). Almost all genes in this set formed a single interconnected network (Supplementary Figure S4), which is particularly notable for the high number of edges converging on several micro-RNA genes (in green). This includes miR-92a, linked to the control of metabolism (Chen & Rosbash, 2017), and miR-92b, implicated in larval locomotion (Chen *et al*., 2012). Even though micro-RNAs tend to have higher number of known interactions, the number of interactions of our candidate micro-RNAs with other candidate genes is in excess of what would be expected at random (*P* = 0.026). We explore some specific links between genes with candidate SNPs and genes that differ in expression in the Discussion.

### Shared candidates for tolerance to larval malnutrition tolerance and adult starvation

In the Introduction we hypothesized that adaptation to chronic juvenile malnutrition would rely on different mechanisms than resistance to acute adult starvation. To address this hypothesis we assessed the overlap between our candidate genes and candidate genes identified in two experimental evolution studies of adult starvation resistance (Hardy *et al*., 2018; Michalak *et al*., 2018). We found that 70 (9%) of our candidate genes were also identified as candidates in at least one of those two studies; the shared genes constituted 10% of candidate genes found by Hardy et al. (2018) and 16% of those found by Michalak et al. (2018), roughly three times more than expected by chance (Figure 5A; Supplementary Table S5). Because all these studies likely missed some relevant genes, and it suffices for a “true” candidate to be a false negative in one of them to be excluded from the overlap, the above numbers likely underestimate the true degree to which the evolutionary response in those studies involved shared genes. For comparison, only 53 candidate genes were shared between Hardy et al. (2018) and Michalak et al. (2018) (14% and 19 % of each other’s pool of candidates), even though both selected for the same trait using similar protocols (Supplementary Table S5). Thus, these results suggest that, contrary to our prediction, evolutionary adaptation to chronic larval malnutrition acted in part on the same genes as the response to selection for adult starvation resistance.

**Figure 5.**
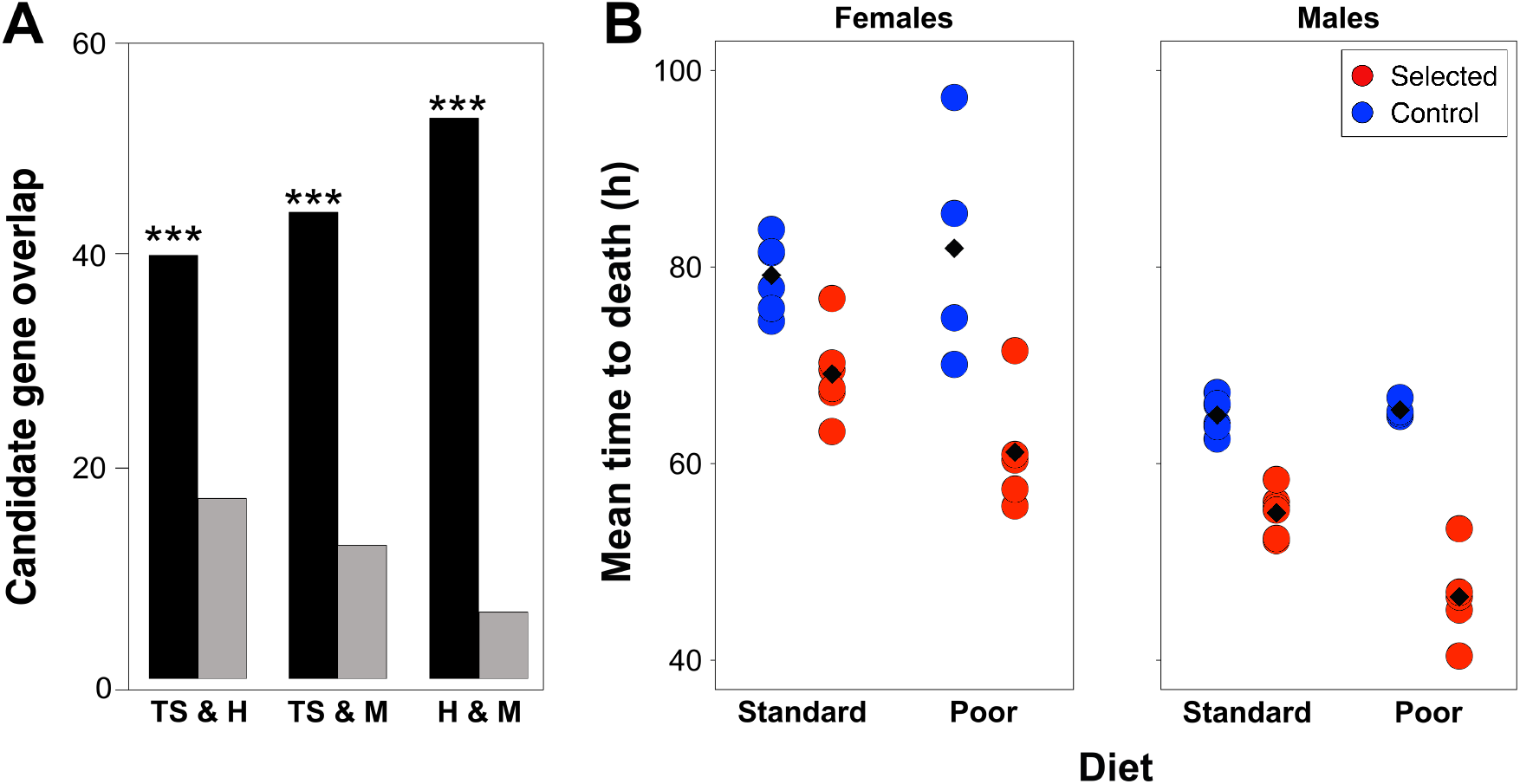
Candidate genes shared with starvation resistance experiments, but reduced starvation resistance in Selected populations. (A) Observed (black bars) and expected (grey bars) overlap of candidate genes between this study (TS) and two studies of selection for starvation resistance, Hardy *et al.* 2018 (H) and Michalak *et al.* 2017 (M). Asterisks indicate significantly larger overlap (*P* < 0.001) than expected by chance. (B) Starvation resistance of female and male adult flies from the Selected (red) and Control (blue) populations. Starvation resistance was quantified as time to death on non-nutritional agar, and was assessed in 1-day old flies raised either on standard or poor diet. Each point shows the mean of a specific replicate population; the black diamond is the overall regime mean.

### Populations adapted to larval malnutrition are less resistant to adult starvation

The overlap of candidate genes between our study and those of Hardy *et al.* (2018) and Michalak *et al.* (2018) suggested that adaptation to chronic larval malnutrition shared to some degree genetic architecture, and thus physiological mechanisms, with resistance to acute adult starvation. If so, our Selected populations should show improved adult starvation resistance compared to Controls. To test this prediction, we quantified starvation resistance of males and females, both when raised on standard and poor diet. Contrary to the prediction, Selected flies showed a lower starvation resistance (died sooner when deprived of food) than Control flies (Figure 5B; *F*_1,15.0_ = 74.5, *P* < 0.0001). This held for both sexes (all interactions involving sex *P* > 0.5). The difference was greater in magnitude in flies raised on the poor diet (regime × diet interaction *F*_1,13.2_ = 9.1, *P* = 0.0099); the Selected flies died about 10 h (14 %) earlier than Controls when raised on the standard diet (*t*_17_ = 4.6, *P* = 0.0003) and about 20 h (27%) earlier when raised on the poor diet (*t*_17_ = 7.5, *P* < 0.0001).

These results suggest that experimental evolution of improved tolerance to larval malnutrition traded off with poorer resistance to adult starvation. This may suggest that the overlap of candidate genes for adaptation to larval malnutrition versus adult starvation reflects antagonistic phenotypic changes favored by selection driven by these two types of nutritional stress.

## Discussion

### Adaptation to chronic malnutrition is highly polygenic and mainly regulatory

Our results imply that experimental adaptation to an extremely nutrient-poor larval diet was highly polygenic. We identified >3,000 candidate SNPs, spread across the genome and annotated to >700 genes, that were significantly divergent in allele frequency between the poor diet-adapted Selected populations and the Controls evolved on a standard diet. Many of those candidate SNPs formed linked clusters with highly correlated allele frequencies, as observed in other evolve and resequence studies (Franssen *et al*., 2017; Michalak *et al*., 2018; Barghi & Schlötterer, 2019; Kelly & Hughes, 2019). This indicates that most candidate SNPs were not targets of selection but diverged by genetic hitchhiking. Nonetheless, we identified 131 clusters of candidate SNPs that appeared to evolve independently from one another, suggesting at least as many different targets of selection. These clusters were spread across all chromosomal arms, including several small and two large clusters on the X chromosome. This corroborates previous results from line cross analyses, which indicated large contributions of the X chromosome to improved performance of Selected populations on the poor diet (Vijendravarma & Kawecki, 2013, 2015). The polygenic response to selection is consistent with adaptation to poor diet impinging upon many aspects of physiology, behavior, life history and possibly even morphology (see below).

Genetic hitchhiking makes it inherently difficult to identify the causative genomic targets of selection (Nuzhdin & Turner, 2013). Therefore, following most other studies (Michalak *et al*., 2017; Hardy *et al*., 2018; Hoedjes *et al*., 2019; Kelly & Hughes, 2019), we did not attempt to separate the putative target SNPs from the putative hitchhikers but used all candidates in the downstream analysis. The fact that we find consistent enrichment of certain functional categories of genes, greater connectivity between candidates than expected by chance, and overlap of candidate genes with other studies of nutritional adaptation implies that the ratio of the signal of adaptation to the noise of hitchhiking is sufficient to make biologically interesting conclusions, as discussed below. Furthermore, hitchhiking of non-target alleles is a relevant factor of correlated responses to selection (Falconer & Mackay, 1996) and thus an important mediator of evolutionary trade-offs (Lande, 1982; Sinervo & Svensson, 1998). Finally, such closely linked clusters may represent distinct haplotypes consisting of multiple co-adapted variants. For example, the cluster of candidate SNPs centered at chromosomal position X:14,900,000 and spanning 350 kb, includes several candidate genes with roles in ecdysone and lipid metabolism *(Eo* and its paralogs, *pdgy, Lsd-2* and *dob*). The center of this cluster corresponds to a region that has been subject to a selective sweep during the out-of-Africa expansion of *D. melanogaster* and contains multiple genetic variants unique to temperate populations (Werzner *et al*., 2013), suggesting it might have acted as a “supergene” in adaptation to the new conditions.

### Patterns of allele frequencies hint at balancing selection at some loci

We expected that, after 150 generations of selection, alleles mediating adaptation to the poor diet would reach high frequencies in the Selected populations. This was indeed the case for many of the candidate SNPs. However, we also detected a substantial number of candidate SNPs that were characterized by intermediate allele frequencies in all Selected populations (the “mid-frequency” candidates). Such a pattern could be generated under at least three evolutionary scenarios.

First, these SNPs might represent incomplete selective sweeps in the Selected populations. If so, the fact that they have not reached fixation after 150 generations would imply weak selection and/or low initial frequencies of the “selected” allele.

Second, initially neutral alleles might have segregated at intermediate frequencies in the base population and later increased under directional selection in the Control populations while remaining neutrally polymorphic in the Selected populations. Although the Control populations were maintained on the same diet as the base population, we cannot rule out that the base population had not yet completely exhausted additive variation for fitness on that diet. Furthermore, to synchronize the generations of Control and Selected populations despite a difference in developmental time on the two diets, the Control populations reproduced at a slightly older age than Selected populations (about 9 versus 6 days since eclosion on average) (Kolss *et al*., 2009). While this is well within the flies’ period of maximum reproduction, it might nonetheless have resulted in some selection on delayed reproduction in the Control populations.

Third, stable intermediate allele frequencies may be maintained by balancing selection. Evidence for balancing selection during experimental evolution is relatively scarce (for example, Skrzynecka & Radwan, 2016; Michalak *et al*., 2017) and the underlying evolutionary mechanisms remain elusive. However, several previous studies suggested that high degree of competition due larval crowding induces negatively frequency-dependent selection, possibly mediated by a trade-off between fast resource acquisition and tolerance to accumulation of metabolic waste products (Kojima & Huang, 1972; Gromko & Richmond, 1978; Borash *et al*., 1998). While larval density experienced by the Selected populations was low, the larvae likely experienced resource competition and deteriorating environmental conditions. In contrast to the two above scenarios, balancing selection would act to reduce variation among replicate populations. Consistent with this, among-population variance in allele frequencies at candidate SNPs with intermediate allele frequencies was smaller in the Selected than in the Controls. Together with positive values of Tajima’s *D* (which indicate an excess of intermediate frequency alleles) observed in the Selected populations at the corresponding genomic regions, this provides indirect support for some role of balancing selection in adaptation to the poor diet.

### Genomic differentiation is associated with diverse phenotypic adaptations

One aspect of the evolutionary adaptation of the Selected populations to the poor diet is an improved ability to extract and assimilate nutrients (notably amino-acids and other nitrogenous compounds; Cavigliasso *et al.*, 2020). This has presumably been mediated at least in part by changes in expression of digestive proteases and a higher proteolytic activity in the guts of the Selected larvae (Erkosar *et al*., 2017). Yet, although several candidate SNPs were annotated to upstream regions of digestive enzymes (notably to *Jon66Cii,* which is also overexpressed in the guts of Selected larvae; (Erkosar *et al*., 2017), overall the candidate SNPs show no sign of enrichment in genes involved in digestion.

Rather, the genomic differentiation points to wide-ranging changes in metabolism. This includes an enrichment of candidate SNPs associated with lipid and carbohydrate metabolism. An earlier gene expression study likewise revealed a strong signal of changes in lipid metabolism in these populations, with many genes showing a reduced expression in Selected larvae relative to Controls (Erkosar *et al*., 2017). In particular, a majority of enzymes directly involved in converting dietary carbohydrates and fatty acids into triglycerides, their storage in lipid droplets and their mobilization (Heier & Kühnlein, 2018) show reduced expression in Selected populations and several are also associated with candidate SNPs (Figure 6). The main transcription factor promoting triglyceride synthesis *(SREBP)* is likewise downregulated while its key activator (*SCAP*) carries a non-synonymous candidate SNP. These genomic and transcriptomic results are consistent with the finding that the Selected larvae assimilate less carbohydrates and accumulate less triglycerides than Controls (Cavigliasso *et al*., 2020). This implies that adaptation of the Selected populations to the poor diet not only targeted nutrient acquisition, but also the way the meager nutrients are used.

**Figure 6.**
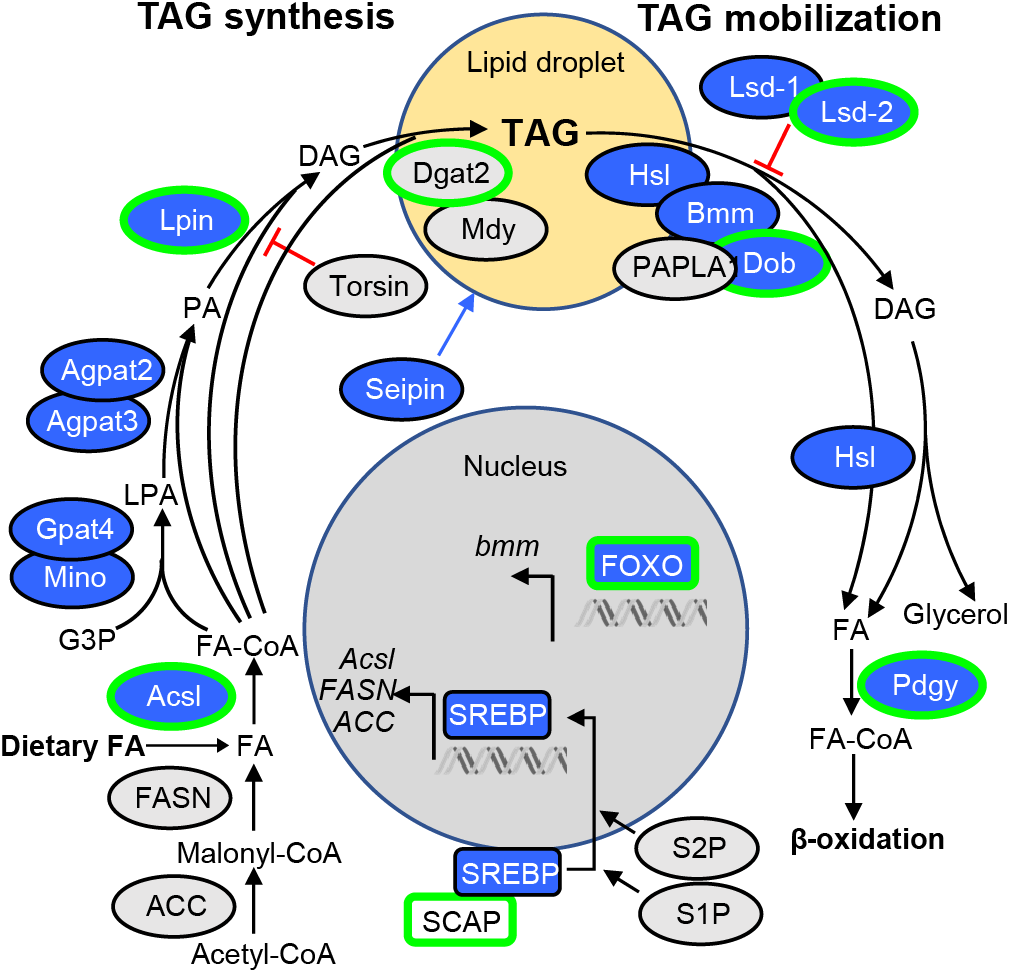
Signature of genomic and gene expression changes in genes involved in triglyceride (TAG) storage and mobilization. Green outline: genes with associated candidate SNPs; blue: genes downregulated in Selected larvae relative to Controls (data from Erkosar *et al*., 2017); no genes are upregulated. Pathway redrawn with modifications from Heier and Kühnlein (2018); FA: fatty acid, G3P: glycerol-3-phosphate, LPA: lysophosphatidic acid, PA: phosphatidic acid, DAG: diacylglycerol.

Analysis of candidate SNPs also indicates a major role of hormonal regulation in the evolutionary differentiation between the Selected and Control populations. This includes cellular response to insulin signaling, which regulates growth in response to nutritional status of the organism. We have previously reported that, when raised on the poor diet, the Selected larvae show a lower expression of *foxo* and many of its target genes than Controls (Erkosar *et al*., 2017). Combining of the expression data from that study with our SNP data suggests involvement of other components of the insulin signaling pathway (Figure 7). Interestingly, while some core members of the pathway tend to be downregulated, several negative regulators are upregulated in the Selected populations. Overall, the changes in insulin signaling appear complex and not interpretable at this stage as a general upregulation or general downregulation of the signaling. Interestingly, with the exception of the fat-body expressed ilp6, neither genomic nor transcriptomic data point to changes in the upstream part of insulin signaling. We also did not find any evidence that the TOR signaling pathway has been a target of adaptation to low nutrient level, despite its key role in regulating the physiological response to nutrients (Templeman & Murphy, 2018). This suggests that the experimental adaptation to chronic larval malnutrition modulated the response to the perceived physiological nutrient level, but not nutrient sensing itself.

**Figure 7.**
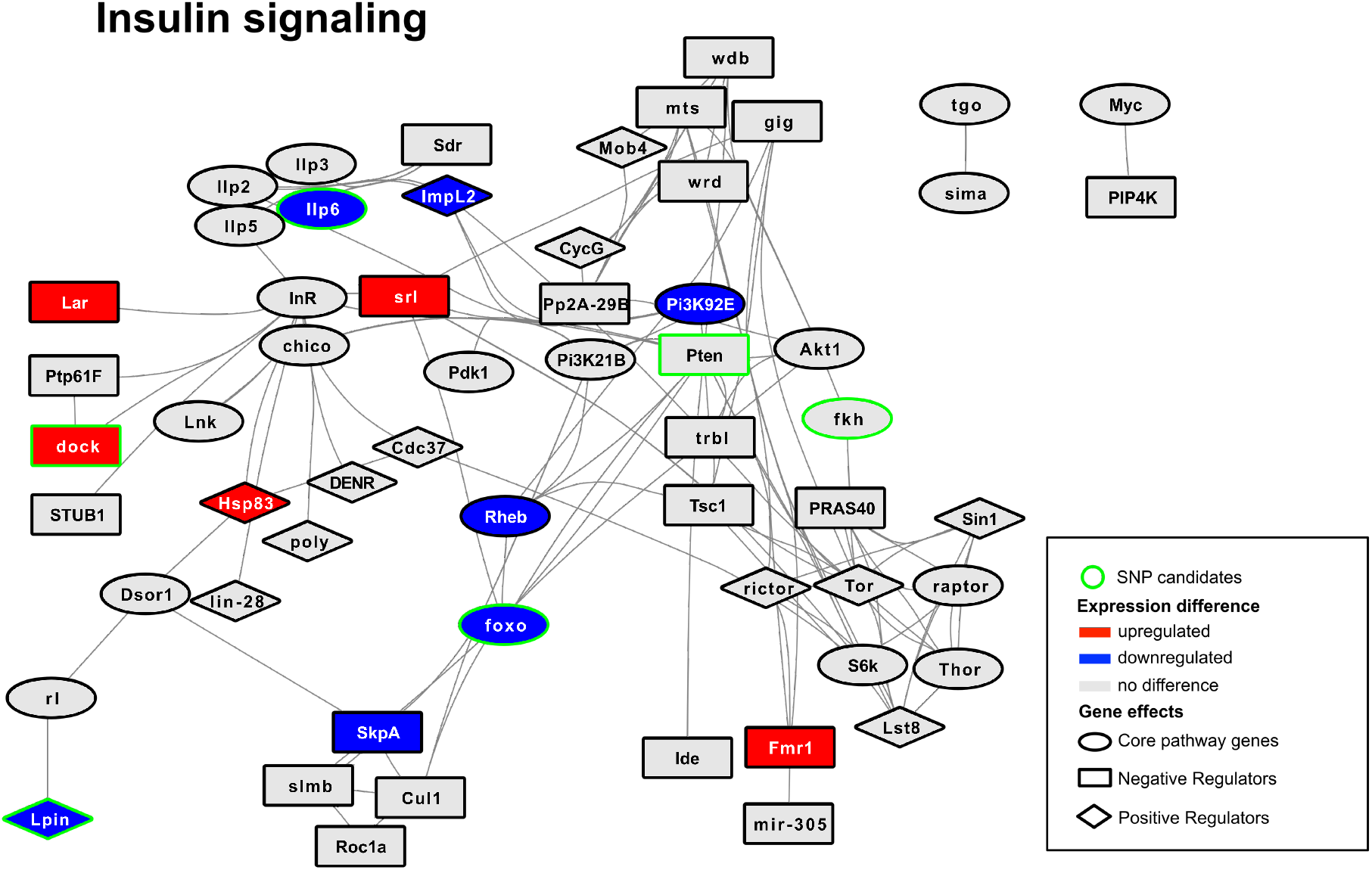
Signature of genomic and expression changes in the insulin signaling pathway. Gene network based on known interactions of genes from the Insulin signaling network obtained from FlyBase. Genes with associated candidate SNPs have a green outline; transcriptomic candidates are highlighted if down-(blue) or upregulated (red) in Selected larvae relative to Controls (data from Erkosar et al. 2017). Symbol shape indicates the role of gene the gene in the pathway (core component, negative regulator, positive regulator), based on information from FlyBase.

We also found a strong signal of genomic changes in ecdysteroid signaling, which regulates molt and metamorphosis. In particular, several genes with known or predicted ecdysone oxidase activity show multiple associated candidate SNPs (including several non-synonymous substitutions) and/or strongly reduced expression. The reaction catalyzed by ecdysone oxidases inactivates ecdysone; thus, lower ecdysone oxidase activity might lead to a faster rise in ecdysone. This finding dovetails with the lower critical weight for metamorphosis initiation and shorter development (independent of diet) evolved by the Selected larvae (Vijendravarma et al. 2012a)

The list of GO terms enriched among candidate genes include a category specific to nervous system: a group of glutamate receptors involved in neuromuscular junctions in the larval body wall, including GluRIIA, which mediates locomotor responses to nutrient shortage in the larvae. The candidates also include multiple genes implicated in larval neuromuscular development, including several genes of the *Beat* family (Li *et al*., 2017) and the micro-RNA miR-92b; deletion of the latter causes larvae to be sluggish (Chen *et al*., 2012). Changes in these genes might have contributed to the reduced locomotion of Selected larvae while foraging (Vijendravarma *et al*., 2012b) and their reduced tendency for tunneling prior to pupation (Narasimha *et al*., 2015). Both might have been favored on the poor diet to save energy.

Finally, the genomic candidates are enriched in genes involved in the adhesion of pupae to surfaces: two salivary gland glue proteins and a protein involved in their secretion. While this may seem idiosyncratic, anecdotical observations suggest that the Selected populations tend to pupate more often close to the surface of the food medium. This strategy may have evolved to save scarce protein. This example underlines the general finding of this study that adaptation to nutrient shortage impinges on manifold and sometimes unexpected aspects of phenotype.

### Adaptation to extreme malnutrition targets some of the same genes as mild variation in nutrition

Hoedjes et al. (2019) have recently analyzed genomic differentiation between *D. melanogaster* populations evolved on larval diets of different nutrient content. However, the range of concentrations of yeast (the main source of most nutrients) used in that study (250, 100 and 25 g per liter) was above those on which not only our Selected populations, but even our Controls evolved (respectively 3.2 and 12.5 g per liter). The difference in the nutritional conditions used in the two studies is also reflected in the egg-to-adult developmental time, which is 11 days on average on the poorest diet used by Hoedjes *et al.* (2019) and May *et al.* (2019) and 14-18 days on our poor diet (Kolss *et al*., 2009). In spite of the milder nutritional conditions and a different founder population in their experiment, 132 (17%) of genes associated with our candidates were also found by Hoedjes et al. (2019), many more than expected by chance. This suggests that at least some aspects of adaptation to the extremely poor larval diet involves in part the same genes that mediate evolutionary response to nutritional conditions well within the normal physiological tolerance of the species.

### Shared genetic architecture points to trade-off, not synergy in resistance to different types of nutritional stress

A major question we asked concerned the relationship between adaptation to chronic larval malnutrition and resistance of adult flies to starvation. As argued in the Introduction, the two main strategies mediating starvation resistance – accumulation of energy reserves and minimizing energy demands while food-deprived (Rion & Kawecki, 2007) – are unlikely to be effective for juveniles facing poor quality diet from birth until maturation. We therefore hypothesized that adaptation to these two types of nutritional stress will largely rely on different genes and molecular mechanisms. Against this expectation, we found a substantial overlap between our genes associated with our candidate SNPs and candidate genes from two genomic studies of starvation resistance (Hardy *et al*., 2018; Michalak *et al*., 2018), suggesting at first sight a shared genetic bases, and thus common molecular and physiological mechanisms.

However, rather than being more resistant, our larval malnutrition-adapted Selected populations turned out to be less resistant to adult starvation than the unselected Control flies. Strikingly, the inferiority of Selected populations to Controls in starvation resistance was greater when the flies were raised on poor diet, even though the Selected populations are adapted to this diet and Controls are not. This suggest that the lower starvation resistance is at least in part a direct consequence of changes in larval development and metabolism that mediate the adaptation of the Selected populations to larval malnutrition, but which affect the phenotype of the emerging adult. In particular, Selected larvae approach pupation with lower lipid stores (Cavigliasso *et al*., 2020), which could directly impact the amount of metabolic reserves of the emerging adults. However, it is also possible that changes in metabolic genes mediating the larval adaptation also impinge on the regulation of metabolic processes in adult flies. Although the main functions of the metabolism of the larvae and adults are different (growth and accumulation of reserves versus conversion of food into locomotion and reproduction), they are largely regulated by the same genes (Murillo-Maldonado & Riesgo-Escovar, 2017). Such non-independence of juvenile and adult metabolism had been proposed to contribute to metabolic disease in humans (Prentice, 2005; Wells, 2007; Vasseur & Quintana-Murci, 2013). This idea is consistent with a strong signal of implication of our candidate genes in catabolic processes; while the amount of energy reserves obviously important for starvation resistance, it is the efficiency of the catabolic processes that will determine how long the animal can survive on those reserves (Rion & Kawecki, 2007).

It has been hypothesized that resistance to different types of environmental stresses may be mediated in large part by the same physiological mechanisms and be positively genetically correlated (Hoffmann & Parsons, 1993; Djawdan *et al*., 1998; Bubliy & Loeschcke, 2005; Sisodia & Singh, 2010). Our results demonstrate that this is not the case even for two types of nutrient shortage, chronic juvenile malnutrition and acute adult starvation. Even though evolutionary responses to these two types of nutritional stress implicate in part the same molecular and physiological processes, they apparently modify them in antagonistic directions. Thus, despite common theme of nutrient shortage and the shared genetic architecture, tolerance to larval malnutrition and to adult starvation are bound by an evolutionary trade-off.

## Materials and Methods

### Experimental Populations

The experimental populations analyzed here originate from an evolution experiment on adaptation to low-nutrient larval diet initiated in 2005. Originally, a base population was derived from several hundred *D. melanogaster* adults collected close to Basel/Switzerland in 1999. Four populations derived from these founders were maintained under standard laboratory conditions at census sizes of >200 individuals for more than 120 generations, allowing them to adapt to the lab conditions. Before the start of the experimental evolution, the four populations were mixed and bred for seven generations at a census size of >1000 individuals to allow recombination to break up large haplotypes in the artificial populations. The evolution experiment was initiated in 2004: six Selected and six Control populations were derived from this base population and maintained ever since at constant laboratory conditions (25°C; 60% humidity) and controlled larval densities (approximately 200 eggs per fly vial) on poor and standard larval diet, respectively. The standard diet consists of 15g agar, 30g sucrose, 60g glucose, 12.5g dry yeast, 50g cornmeal, 0.5g MgSO4, 0.5g CaCl2, 30mL ethanol, 6mL propionic acid, and 1g nipagin per 1L water; the poor diet contains a quarter of the amounts of sugars, yeast, and cornmeal of the standard medium. Under both regimes, adults were transferred to standard diet with supplemental yeast to facilitate oviposition at every generation; the nutritional restriction was thus limited to the larval stage (Kolss *et al*., 2009).

In addition to laboratory natural selection to chronic larval malnutrition we further selected against delayed development, which is a common correlated phenotypic response to malnutrition and has similarly been observed in this selection experiment (Kolss *et al*., 2009). Long development times may be highly maladaptive in nature given that rotting fruit, which represent the natural substrate for developing larvae, decay rapidly and are thus only available for a limited amount of time. We therefore chose flies that developed within 14 days from egg to adult to contribute to the next generation for both regimes. Flies were then maintained for another six days on standard diet before the next generation was initiated.

### DNA extractions and genome sequencing

Genomic DNA was extracted from whole flies for each of the six selected and six control populations in 2014, after approximately 150 generations. We pooled 400 females per population and homogenized them in liquid nitrogen prior to pooled DNA extractions using the Qiagen Dneasy Blood and Tissue DNA extraction kit (Qiagen, Hilden, Germany) with modifications (Kapun *et al*., 2020). DNA of each sample was sheared in a Covaris instrument. Sample-wise library preparation following manufacturer’s instructions for paired-end sequencing were carried out using the Illumina TrueSeq Nano Library kit (Illumina, San Diego, CA, USA). DNA pools were sequenced in two multiplexed batches of 6 samples (3 selected and 3 control population per batch) each on an Illumina 2500 sequencer at the genomics technologies facility of the University of Lausanne, yielding paired-end sequences of 100bp lengths.

### Mapping pipeline and variant calling

Raw FASTQ reads from all libraries were trimmed and filtered using *cutadapt* (v.1.8.3; Martin, 2011) to remove low-quality bases (minimum base PHRED quality = 18; minimum sequence length = 75bp) and Illumina-specific sequencing adaptors. We only retained read pairs where both reads fulfilled all quality criteria and mapped these against a compound reference consisting of the genomes from *D. melanogaster* (v.6.04) and common commensals and pathogens, such as *Saccharomyces cerevisiae* (GCF_000146045.2), *Wolbachia pipientis* (NC_002978.6), *Pseudomonas entomophila* (NC_008027.1), *Commensalibacter intestine* (NZ_AGFR00000000.1), *Acetobacter pomorum* (NZ_AEUP00000000.1), *Gluconobacter morbifer* (NZ_AGQV00000000.1), *Providencia burhodogranariea* (NZ_AKKL00000000.1), *Providencia alcalifaciens* (NZ_AKKM01000049.1), *Providencia rettgeri* (NZ_AJSB00000000.1), *Enterococcus faecalis* (NC_004668.1), *Lactobacillus brevis* (NC_008497.1), and *Lactobacillus plantarum* (NC_004567.2) using *bwa mem* (v.0.7.15; Li, 2013) with default parameters. After that, we removed read duplicates and reads with mapping qualities < 20 using *Picard* (v.1.109; http://picard.sourceforge.net) and re-aligned sequences flanking indels with *GATK* (v.3.4-46; McKenna *et al*., 2010). We then used *samtools mpileup* (v.1.3; Li & Durbin, 2009) to merge quality filtered BAM files from all samples and performed heuristic SNP calling on the mpileup file based the following parameters using *PoolSNP* (Kapun *et al*., 2020): (1) minimum coverage from all samples ≥ 10x, (2) maximum coverage from all samples ≤ the 95^th^ coverage percentile for a given chromosome and sample, (3) minimum read count for a given allele ≥ 20x and (4) minimum read frequency of a given allele ≥ 0.01 across all samples pooled. We further excluded positions where ≥ 20% of all samples did not fulfill the minimum and maximum coverage thresholds, which overlapped with indel polymorphisms or were located within 5 bp distance or which were spanned by known transposable elements (TE) based on the *D. melanogaster* TE library v.6.10.

All sequenced libraries were of high quality with average raw PHRED-scaled base qualities exceeding 28 in all samples. Our mapping pipeline with stringent quality filtering resulted in similar coverages across all major chromosomal arms and yielded sample-wise average read depths ranging between 30x to 46x (see Table S1).

### Multivariate analyses of SNP frequencies

We performed a principal component analysis in SAS v. 9.4 on allele frequencies of 976,247 SNPs which fulfilled the coverage threshold criteria in all twelve replicate samples; because allele frequencies are already measured on the same scale, this was done on the covariance rather than correlation matrix. To test for separation between the evolutionary regimes we compared their scores on the principal component axis with Welch’s *t*-test.

### Population genetic analyses

To characterize the extent and distribution of genetic variation in each sample, we estimated the population genetics statistics *π,* Watterson’s *θ* and Tajima’s *D* focusing on SNPs located on the six major chromosomal arms *(X, 2L, 2R, 3L, 3R* and *4).* Prior to calculations, we homogenized all sequencing data to an even 30-fold coverage to control for the coverage sensitivity of Watterson’s *θ* and Tajima’s *D*. Sites with coverages greater than 30-fold were randomly subsampled without replacement. In contrast, we thirty times randomly sampled reads with replacement for sites where sequencing depths were below the target coverage (Kofler *et al*., 2011; Kapun *et al*., 2020). For each statistic, we calculated genome-wide and window-wise averages in sliding windows of 200 kb length with 200 kb step-size with *PoolGen* (Kapun *et al*., 2020), which employs statistical corrections for population genetics estimators as described in Kofler *et al.* (2011). These corrections account for sampling biases and sequencing errors as a result of pooled re-sequencing (Futschik & Schlötterer, 2010).

We estimated the effective population sizes *(N_e_)* of the Selected and Control populations based on among-populations variance in allele frequencies at putatively neutral SNPs. Under drift, the expected variance in allele frequencies at a SNP locus among replicate populations as estimated from Pool-seq data is

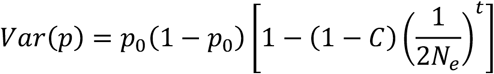

where *p*_0_ is the initial allele frequency, *t* = 150 is the number of generations since the populations were established from a common base, and *C* is a correction factor to account for variance due to genetic drift and sequencing (Jonas *et al*., 2016). Estimating *p*_0_, Var(*p*) and *C* and solving the equation yields a SNP-specific estimate of *N_e_*. Following Jónás et al. (2016), for each population *j* we first calculated *C_j_* = 1/2*S* + *1/R_j_* + 1/(*SR_j_*) where *R_j_* is the number of reads used to estimate the allele frequency (i.e., the read depth) of population *j,* and *S* is the number of individuals included in the pooled sample (400 for all populations). *C* for a given SNP was then calculated as the arithmetic mean of *C_j_* across populations (*C* is thus a function of harmonic mean of read depth across populations). Because sequence data from the initial base population were not available, the initial frequency *p*_0_ for each SNP was estimated as the average frequency of this SNP across all 12 populations at time *t* (Hardy et al. 2018). Because we wanted to compare *N_e_* estimates between the evolutionary regimes, we obtained separate estimates of allele frequency variance for the Selected populations as

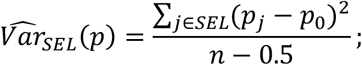

where *n* = 6 is the number of replicate populations per regime; an analogous equation was used to obtain 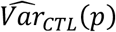 for the Control populations. The subtraction of 0.5 from the denominator reflects the estimation of *p*_0_ using one degree of freedom, half of it contributed by data from each regime. (Note that the average of 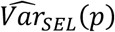 and 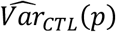 equals the variance of *p_j_* across all 12 populations.)

We applied the above approach to SNPs that were in short introns (<60bp length), and thus thought to be neutral (Parsch *et al*., 2010; Clemente & Vogl, 2012). Of those, we only used SNPs with minimum coverage of 10 in each population and mean allele frequencies between 0.4 and 0.6 (724 SNPs on chromosomes 2 and 3, 103 SNPs on the X); *N_e_* estimates based on SNPs with asymmetric mean allele frequency tend to be biased upward (Hardy *et al*., 2018). For each of these SNPs we obtained a separate *N_e_* estimate for the Selected and Control populations; we took the median of these values (separately for the autosomes and the X) as our final estimates. The significance of differences in *N_e_* estimates between regimes was assessed by randomization. We generated 100 data sets by dividing the 12 populations at random in two groups each consisting of three Control and three Selected populations. We then estimated *N_e_* for the two groups for each of these randomized data and used the distribution of pairwise differences of these estimates as a null distribution to which we compared the difference of *N_e_* estimates between Selected and Control populations.

### Detecting candidates SNPs

While some recent evolve and resequence studies took advantage of temporal trajectories of alleles from the beginning of experimental evolution, such data are not available for our populations, limiting us to the analysis based on the comparison of allele frequencies between sets of populations exposed to different regimes. Because the expected distribution of SNP allele frequencies under a combination of selection and drift does not correspond to the assumptions of any classic statistical tests, a range of heuristic statistical approaches has been used to identify putative candidate SNPs in this type of data. Some studies employed approaches that emphasize the consistency of allele frequency differentiation among replicate populations, such as the “diffstat” statistics (Turner *et al*., 2011) or generalized linear mixed models (GLMM, e.g. Jha *et al*., 2015a) or a general linear model (e.g., Hoedjes *et al*., 2019). Other studies relied on test that emphasize the mean difference in allele frequency and are insensitive to heterogeneity among replicates, such as Fisher’s exact test on pooled replicate populations (e.g., Burke *et al*., 2011) or Cochran-Mantel-Haenszel (CMH) test (e.g., Orozco-terWengel *et al*., 2012; Michalak *et al*., 2018). (It should be noted that even though the CMH test uses stratified data (e.g., pairs of populations), it assumes that the odds ratios are homogenous across strata and the value of the CMH statistics is little affected by heterogeneity (Wiberg *et al*., 2017).) This second approach is justified because an initially rare SNP allele is likely to become lost from some replicate populations due to drift even if it has a strong selective advantage. This will generate a highly heterogenous pattern where the allele is at high frequency in most selected populations but absent in one or a few; this pattern can be detected by an approach that is insensitive to heterogeneity, as we demonstrate in this paper.

We employed a two-pronged approach to detect adaptive alleles showing either of the above patterns. (1) We used SNP-wise Fisher’s Exact tests (FET), which targeted SNPs that were on average strongly differentiated between the regime groups, but not necessarily consistent across replicate populations (Remolina *et al*., 2012). To equalize contribution of each population we subsampled each SNP position to a 30-fold coverage in each library as described above and then we pooled SNP allele counts from all populations within each regime. The resulting 2×2 contingency tables were tested with FET in *R* for each SNP separately. (2) Independently, we fitted SNP-wise generalized linear mixed models (GLMM) with a binomial error structure to allele counts without sub-sampling, which detect SNPs that might have only moderately diverged in frequency between the regimes, but in a way that is consistent among the replicate populations. We used the *glmer* function of the *R* package *lme4* (Bates *et al*., 2015) to calculate GLMMs of the form: *y*_i_ = *Regime* + *Population(Regime)* +*ε*_i_;, where *y*_i_ is the allele frequency of the *i*^th^ SNP, *Regime* (control versus selected) is a fixed factor, *Population* is a random factor for replicate populations nested within *Regime* (to account for overdispersion) and *ε*_i_ denotes a binomial error term. We tested for significant effects of the evolutionary regime by comparing the full model to a reduced model without the fixed effect by means of likelihood ratio tests using the *R* function *anova.*

As has been noted previously, FET applied to pooled data results in a large excess of small *P*-values (Turner *et al*., 2011), and even for GLMM the null distribution of *P*-values is not uniform (Jha *et al*., 2015b), i.e., there are not true *P*-values, but rather should be treated as statistics with an a piori unknown distribution. To account for this, we followed the approach of Jha *et al.* (2015b): we estimated FDR by comparing the distribution of *P*-values obtained in each of the two above-mentioned statistical tests with a corresponding empirically estimated null distribution. To generate these null distributions of *P*-values, we permuted the input data by randomly assigning three Selected and three Control samples to each of two treatment groups and performed the FET and GLMM as described above. This was done on five independently permuted data sets. From these permutations we calculated, for any cutoff value of 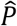, the expected (null) cumulative distribution function 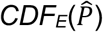 as the proportion of P-values 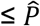 (averaged over the five permutations). The observed cumulative distribution function 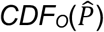 was analogously defined for the original (non-permuted) data. FDR (i.e., the adjusted *P*-value) corresponding to a given raw *P*-value *P* was then estimated as

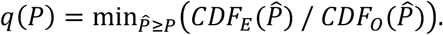

Because the null distributions of raw *P*-values were different between the two tests, we performed this procedure separately for the FET and GLMM. Sites with *q* ≤ 0.05 in at least one of the two tests were considered significant. Prior to all subsequent analyses, we further filtered the resulting candidate SNP dataset by removing those where average allele frequency difference between Selected and Control populations was smaller than an arbitrary threshold of 0.3. This removed 101 SNPs detected by GLMM, that had very uniform allele frequencies across replicate populations, not expected after 150 generations at a rather small population size. Even if these SNPs were under selection, the small difference in mean allele frequency despite 150 generations of selection would imply a very small selection coefficient.

### Association among candidate SNPs and genes and the impact of variable recombination rates

To assess the extent of linkage disequilibrium resulting in non-independent evolution of linked SNPs we calculated within-regime correlations between allele frequencies at different candidate SNPs. To this end, we first expressed allele frequencies as deviations from the regime means (residuals), thus removing the component of covariance which is due to evolution under the same selection regime. Then we calculated pairwise correlations of these frequency deviations between all pairs of SNPs on the same chromosomal arm across all 12 populations. The fact that allele frequencies at two neighboring SNPs have highly correlated residuals is most parsimoniously explained by a combination of linkage disequilibrium and drift (including founder effect). These correlations were plotted as a heat map.

Complementary to the analyses of pairwise correlations of allele frequencies, we investigated and similarly visualized pairwise recombination rates among candidates. We therefore downloaded average recombination rate estimates in 100-kb windows (Comeron *et al*., 2012) from the *Drosophila* recombination rate calculator (https://petrov.stanford.edu/cgi-bin/recombination-rates_updateR5.pl). Then, we converted genomic positions to match the *Drosophila* reference v.6 coordinates and calculated approximate pairwise recombination probabilities by summing site-specific recombination rate estimates between candidate SNPs on the same chromosomal arm.

### Functional characterization of candidate SNPs

Based on the Ensembl genome annotation (v. BDGP6.82) we used SNPeff (v.4.2; Cingolani *et al*., 2012) with standard parameters to annotate the total SNP dataset for genomic features and to predict genetic effects. We further identified SNPs located in regulatory elements using information for *cis*-regulatory modules (CRMs) and Transcription factor binding sites (TFBS) from the REDfly database (v.5.2.2; Gallo *et al*., 2006). We employed Fisher’s exact test s with Bonferroni correction (corrected *α* =0.006) to test whether SNPs mapping to different genomic features were under-or overrepresented among candidate datasets relative to all other polymorphic sites.

Using *Gowinda* (Kofler & Schlötterer, 2012), which performs permutation tests based on randomly drawn SNPs without replacement, we further tested for enrichment of gene ontologies (GO) in all candidates jointly as well as in candidates for balancing and directional selection separately. *Gowinda* allows to control for a spurious overrepresentation of GO terms composed of long genes, since these have a higher probability to contain falsepositive candidate SNPs compared to short genes.

We further tested for functional links between our list of genomic candidates and previously published transcriptomic data from the same experimental populations (Erkosar *et al*., 2017). We therefore focused on genes that were previously identified as candidates for differential expression with respect to selection regime based on *limma voom* (Law *et al*., 2014) analyses as described in Erkosar et al. (2017). We used the *R* package *SuperExactTest* (Wang *et al*., 2015), which estimates predicted intersections among the datasets based on the size of a common statistical background population. This background was calculated based on all genes that harbored polymorphic SNPs in the genomic datasets and that had ≥ 1 count per million in at least 6 samples of the RNA-Seq data. We further used information about known interactions among genes from the DROID database (Yu *et al*., 2008; Murali *et al*., 2011) to identify functional links between candidates in the joint genomic and transcriptomic datasets. The aim of this approach was to trace for allelic changes that may lead to variation in expression patterns in functionally connected genes such as transcription factors and their target genes. To this end we further used a FlyBase database (“gene_group_data_fb_2019_02.tsv”) to roughly classify candidate genes into (1) transcription factors, (2) enzymes, (3) signaling genes, (4) channel proteins, (5) microRNA’s and (6) unclassified genes. We further used a FlyBase genome annotation file (“dmel-all-r6.17.gtf.gz”) to categorize candidates by gene-length. Based on these classifications, we 1000 times randomly drew non-candidate gene sets from the gene background that matched in classification and length to the true candidate lists and counted the number of interactions among genes within each random gene set. Based on the distribution of numbers of interactions in the 1000 random datasets, we calculated an empirical p-value using the number of interactions of the true candidate set as our threshold. Using *cytoscape* (Shannon *et al*., 2003), we then visualized the interaction networks and highlighted the gene classification by color and the source of each candidate gene (Genomic and/or Transcriptomic candidate) by the shape of the network edges.

At last, we tested to which extent the complete list of candidate genes, as well as separately the “mid-frequency “ and “high frequency” subsets, are shared with Hoedjes the studies of Hardy et al. (2018) and Michalak et al. (2018), which both investigated the adaptation for starvation resistance during experimental evolution. For each combination of datasets, we calculated expected and observed gene overlap and tested for significance using the *R* package *SuperExactTest* as described above.

### Assay of starvation resistance

We measured starvation resistance in flies of the Selected and Control populations raised as larvae on the standard or the poor diet. This took place after 219 generations of selection, followed by three generations on the standard diet to remove the effect of parental nutritional environment (Vijendravarma & Kawecki 2010). Flies of both sexes were collected within 24 h of emergence and transferred in groups of 15 to vials with 3.3% agarose (three replicates per population and sex). Dead flies were counted every 8-10 hours; flies that died between two checks were assigned the middle of the interval as their time to death. Mean time to death was calculated for each replicate and analyzed as the response variable in a general mixed model, with regime, diet and sex as fixed factors, and population (nested in regime) and its interactions with diet and sex as random factors using the R packages lme4 (Bates *et al*., 2015) and emmeans (Lenth *et al*., 2019). For populations C2, C4 and S4 only flies raised on standard diet were assayed; the corresponding poor diet culture bottles did not provide sufficient numbers of adults.

## Acknowledgements

We thank T. Flatt and for helpful discussion and general support with the project. We also acknowledge the support of the Department of Ecology and Evolution and the Vital-IT bioinformatics core facility at the University of Lausanne. We also thank L. Savary for helping with experiments. All sequencing was carried out by the Genomic Technologies Facility at the Center for Integrative Genomics of the University of Lausanne. TJK was supported by the Swiss National Science Foundation (grant number 31003A_162732to). The work of MK was funded by the Swiss National Science Foundation (grant number PP00P3_133641 Thomas Flatt) and supported by the Austrian Science Fund (grant number FWF P32275 to MK). BH’s work was supported by an Ambizione fellowship of the Swiss National Science Foundation (PZ00P3_161430).

## Author Contributions

TJK, RCS and MK designed the study. TJK, RCS and MK performed bioinformatic analyses. CD performed phenotypic assays. TJK, BH, BE, CD and MK contributed to the interpretation and analysis of the data. TJK and MK wrote the manuscript with the critical input of BH, BE and CD.

## Data Accessibility

The unpublished raw sequence data used in this study are available from NCBI SRA (BioProject PRJNA515033). The raw starvation phenotype data are provided in the supplementary material. Novel bioinformatics tools developed for this project and a description of the bioinformatic analysis pipeline can be found at https://github.com/capoony/GenomicsOfLarvalMalnutrition

