## Supplementary material for "The genomic architecture of adaptation to larval malnutrition points to a trade-off with adult starvation resistance in *Drosophila*": SI Text

### Supplementary analyses

#### Population genetic analyses

Chromosome-wide patterns of within-population genetic variation were similar between the regimes. We found no differences between Selected and Control populations for genome-wide estimates of $\pi$ and Watterson's $\theta$ on any autosomal arm (Mann Whitney U tests; all $W < 25$, $P > 0.05$). There was a marginally significant deviation on the *X* for $\pi$ (Mann Whitney U tests; $W = 31$, $P = 0.041$), but not for $\theta$. These results are complementary to the estimates of $N_e$ and indicate overall similar effective population sizes in both regimes. Chromosome-wide $\pi$ values ranged from 0.0004 to 0.0012 for the *X* and 0.0021 to 0.0025 for autosomes respectively (see Supplementary Table S1) which is markedly lower than previous estimates from other natural and experimental *D. melanogaster* populations (see Figure S1) based on pooled sequencing data. Our experimental design that allowed only 200 adults per replicate populations to contribute to next generation and the long course of the experiment in comparison to other experimental evolution studies may have resulted in loss of genetic variation due to drift. When investigating window-wise chromosomal patterns of genetic variation, we identified several genomic regions with statistically significant differences in $\pi$ and $\theta$ with respect to regimes (see Figure S1). Curiously, we found chromosomal regions with strong reductions of genetic variation not only in Selected, but also in consistent across Control populations (see Figure S1).

#### Chromosomal inversions

We tested to which extent genetic diversity could be influenced by chromosomal inversions, which have been shown to have a strong impact on recombination rates and genetic variation (Kapun and Flatt 2018). Based on inversion-specific marker SNPs, the cosmopolitan inversions *In(2L)t, In(2R)NS, In(3L)P, In(3R)C, In(3R)Mo, In(3R)K* and *In(3R)Payne* were completely absent or segregated at very low frequencies (< 5%) in all populations. These findings may either indicate that inversion were at low frequencies or completely absent at the beginning of the selection experiment or alternatively decreased in frequency during the course of the experiment. Such a pattern would be consistent with similar findings from other lab-based experimental evolution studies, which all report that common cosmopolitan inversion rapidly decrease in frequency when maintained under laboratory conditions (Kapun et al. 2014). We therefore conclude that inversions only played a limited role during the evolutionary process in our experiment.

### Supplementary Figures

#### Figure S1

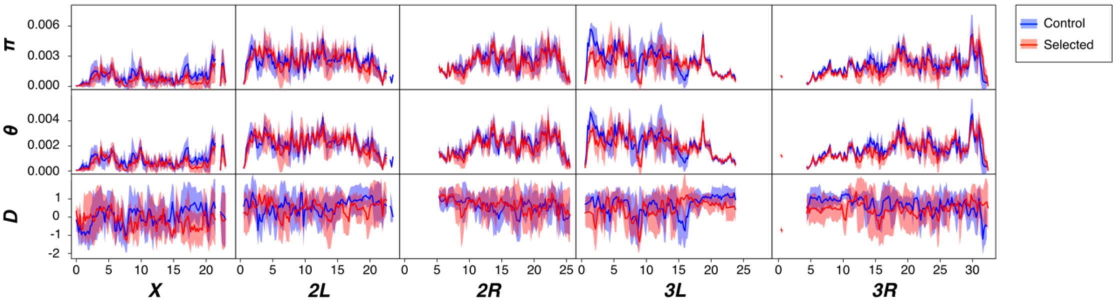


**Supplementary Figure S1: Population genetic analysis of candidates for selection.** The panels show chromosome-wise averages for the population genetic estimators *π*, *θ* and Tajima's *D* in non-overlapping windows of 200 kbp size. Similar to Figure 2, solid lines and semi-transparent polygons show means and standard deviations for these estimators, respectively, that were calculated from six selected (red) and six control (blue) replicate populations.

**Supplementary Figure S2**

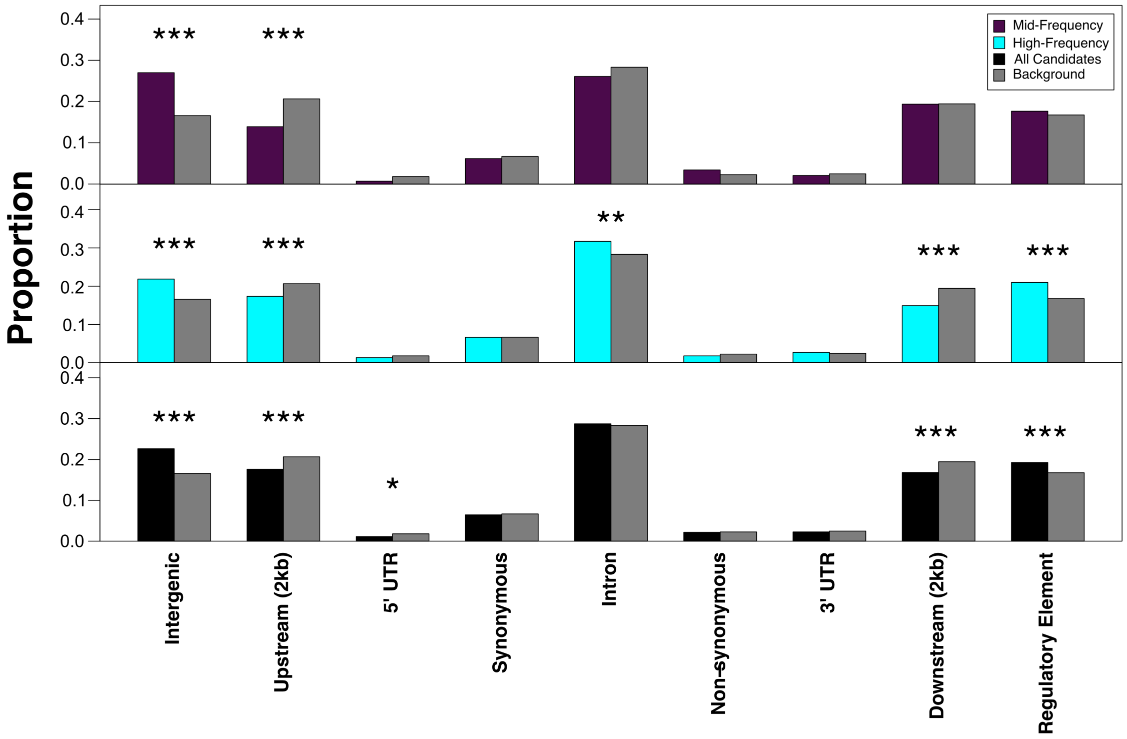


**Supplementary Figure S2. Over- and underrepresentation of candidate SNPs in genomic functional classes.** Histograms showing the proportion of candidates for balancing selection (in orange), directional selection (in cyan) and for all candidates (in black) in functional SNP effect classes, which were assessed by SNPeff annotation and based on the Redfly database. Expected proportions were estimated from all non-candidate SNPs and are highlighted in grey. Significant over- or underrepresentation of selected SNPs was assessed by chi-square tests with Bonferroni corrections (corrected $\alpha$ =0.006) comparing the counts of candidate and non-candidate SNPs in a given feature against the remaining candidate and non-candidate SNPs. ** $P < 0.01$; *** $P < 0.001$

**Supplementary Figure S3**

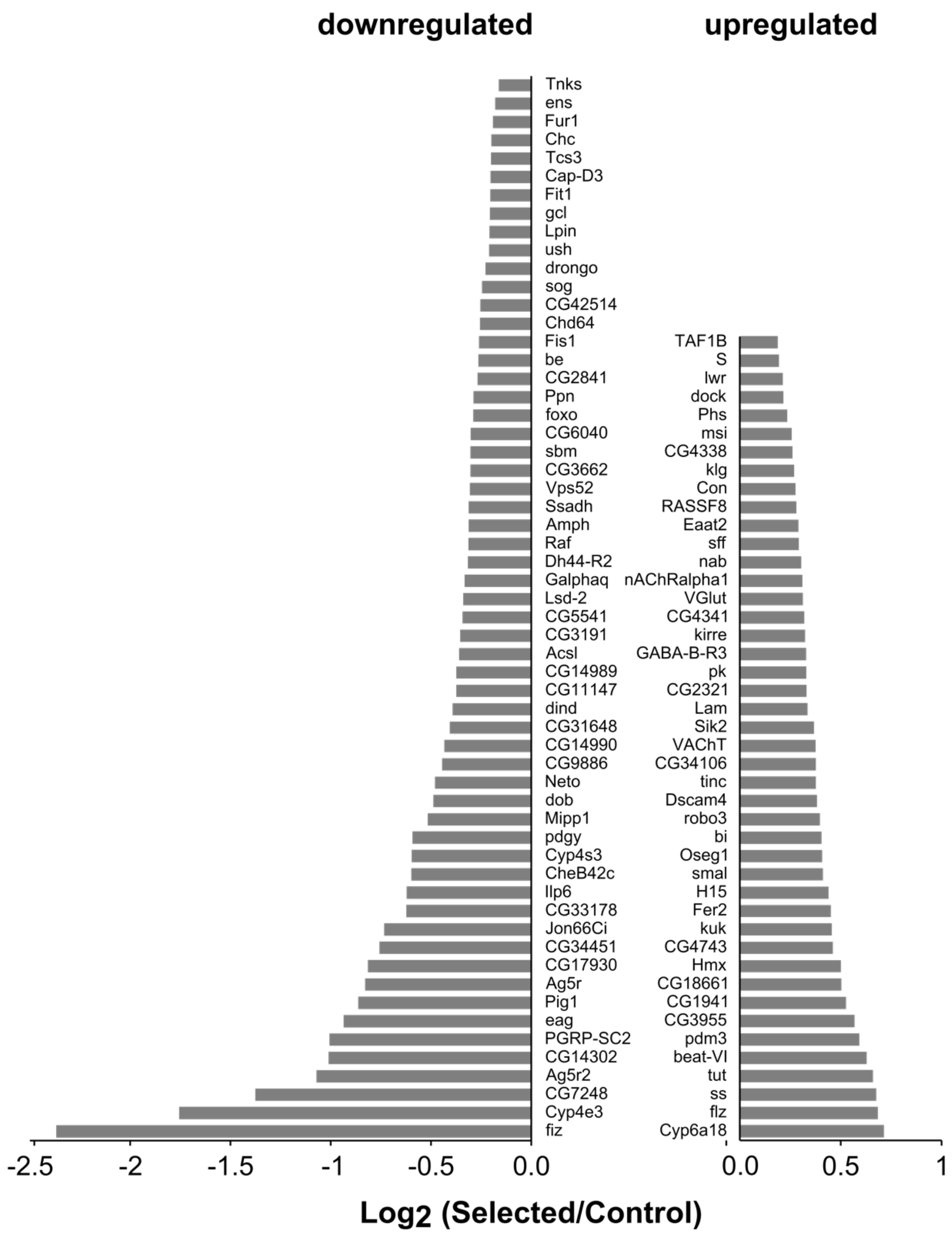


**Supplementary Figure S3**. **Expression patterns of candidate genes shared across genomic and transcriptomic analyses**. Barplots showing the $log_2$-fold differential expression between Selected and Control populations for genes which have been identified as candidates both in the genomic analyses of this study and transcriptomic analyses in Erkosar et al. (2017). Left bars show downregulated and the right bars upregulated candidate genes.

**Figure S4**

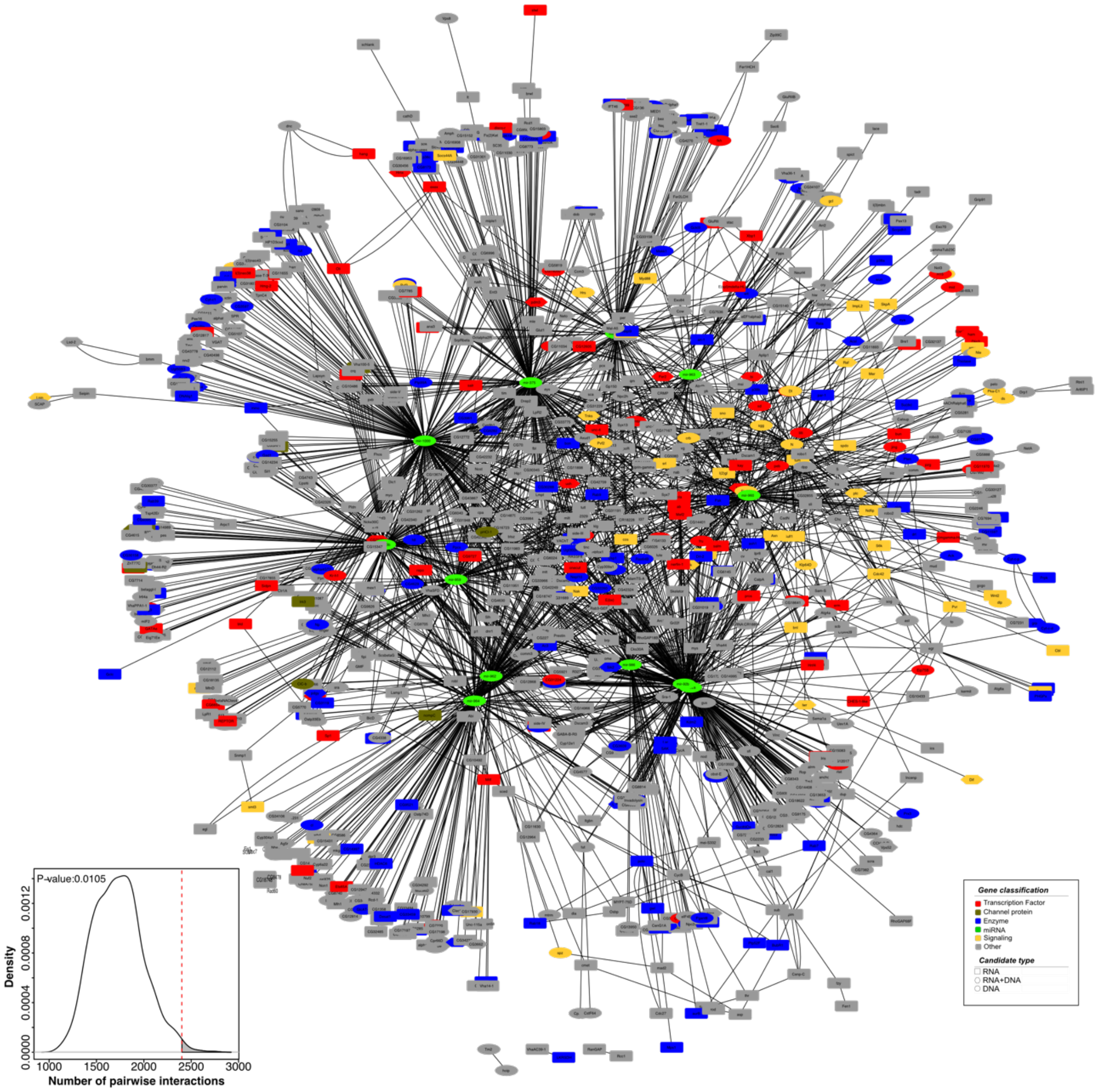


**Supplementary Figure S4: Interaction among all candidate genes.** A gene network showing interactions among all genomics candidates and genes which show significant differential expression among selection regimes (Erkosar et al. 2017). Colors depict a rough classification of genes according to gene function and symbol shape highlight the data source of each candidate. The subplot at the bottom left depicts the distribution of numbers of pairwise interaction in 1000 randomly drawn non-candidate gene sets, that matched the true dataset in number of candidates, gene lengths and gene classifications. The vertical red line highlights the number of interactions in the “true” datasets and was used as a threshold to calculate empirical p-values that are shown in the top-right corner.
